# Enhanced biodegradation of naphthalene by *Pseudomonas* sp. consortium immobilized in calcium alginate beads

**DOI:** 10.1101/631135

**Authors:** Kunal Dutta, Sergey Shityakov, Ibrahim Khalifa, Saroj Ballav, Debarati Jana, Tuhin Manna, Monalisha Karmakar, Priyanka Raul, Kartik Chandra Guchhait, Chandradipa Ghosh

**Author notes:** Corresponding author:* Prof. Chandradipa Ghosh, Microbiology and Immunology Laboratory, Department of Human Physiology with Community Health, Vidyasagar University, Midnapore - 721102, West Bengal, INDIA, /555/557 Extn. 450.

## Abstract

Polycyclic aromatic hydrocarbons (PAHs) belong to a large group of organic pollutant which considers as a potential health hazard to living beings. Herein, naphthalene biodegradation potential by free and immobilized *Pseudomonas putida* strain KD10 and *Pseudomonas* sp. consortium were studied. Additionally, naphthalene 1, 2-dioxygenase (*nah*Ac) was sequenced and analyzed, which reveals two altered amino acid residues. However, the altered amino acid residues are not present in the vicinity of the active site. The gas-phase binding free energy (ΔG_London_) of the mutant variant of naphthalene 1, 2-dioxygenase was -7.10 kcal mol^-1^ which closely resembles the wild type variant. Naphthalene biodegradation rate by *Pseudomonas putida* strain KD10 was 79.12 mg L^-1^ day^-1^ and it was significantly elevated up to 123 mg L^-1^ day^-1^ by the immobilized *Pseudomonas* sp. consortium. The half-life (*t_1/2_*) for naphthalene biodegradation was 3.1 days with the inhibition constant (*k_i_*), substrate saturation constant (*k_s_*) and maximum specific degradation rate constant (*q_max_*) of 1268 mg L^-1^, 395.5 mg L^-1^ and 0.65 h^-1^, respectively, for the *Pseudomonas putida* strain KD10. However, the t*_1/2_* value was significantly reduced to 2 days along with *k_i_*, *k_s_* and *q_max_*values of 1475 mg L^-1^, 298.8 mg L^-1^ and 0.71 h^-1^, respectively, by the immobilized *Pseudomonas* sp. consortium. The GC-MS data suggest that KD10 might follow D-gluconic acid mediated meta-cleavage pathway of catechol biodegradation. It is concluded that naphthalene biodegradation performance by immobilized *Pseudomonas* sp. consortium was superior to free or immobilized *Pseudomonas putida* KD10. Microbial consortium immobilization could be a useful tool for water quality management and environmental remediation.

**Highlights:**

- Superior naphthalene biodegradation by *Pseudomonas* sp. consortium immobilized in calcium alginate beads.
- A common mutation prone amino acid stretch inside chain A of naphthalene 1, 2-dioxygenase has been identified.
- A new naphthalene biodegradation pathway by *Pseudomonas putida* strain KD10 has been proposed.

## 1. Introduction

Polycyclic aromatic hydrocarbons (PAHs) are considered as the potential health hazard for living beings (Kumari et al., 2018). Environmental agencies including US-EPA, European Union, Environment-Canada, registered PAH as the priority pollutants that require immediate human intervention (Wang et al., 2018). The physicochemical properties of PAH make them a major contributor for soil and groundwater contamination through bio-magnification (Norris, 2017). Moreover, according to the Environmental Health Hazard Assessment, U.S.A., naphthalene (NAP) concentration beyond 170 ppb is not safe drinking (Bruce et al., 1998). Different orthodox and expensive techniques for environmental remediation, *viz.*, incineration, gasification, plasma-gasification have been replaced by green-technologies such as bioremediation, phytoremediation, nanoremediation, *etc.* (Thomé et al., 2018). Bioremediation is considered as the most cost-effective and eco-friendly oil spill management technique (Wilson and Jones, 1993). However, while considering the vast volume of a mobile open water system, bioremediation stumble upon several limiting factors such as the low local concentration of the effective microorganisms, loss of active microorganisms, *etc.* (Chen et al., 2017). Conversely, cell immobilization provides several advantages such as, it helps to retain high local concentration of the effective microorganisms, keep intact bacterial cell membrane stability and can be stored for future reuse (Bhardwaj et al., 2000). Overall, cell immobilization can markedly improve the stability and efficiency of the bioremediation process (Mrozik and Piotrowska-Seget, 2010; Tyagi et al., 2011). Cell immobilization using calcium alginate beads (CABs) is a convenient option where maximum cells remain viable and can tolerate high concentration of the toxicant (Lee and Heo, 2000). Moreover, calcium alginate is nontoxic to the bacterial cell and it has low production cost which facilities easy reuse (Bhardwaj et al., 2000). Biodegradation of a toxicant (complex nutrient for bacteria) by the microbial consortium can efficiently enhance biodegradation rate (Kumari et al., 2018). This rate enhancement could be achieved by several ways such as, different biodegradation pathways of each individual bacterium (Dutta et al., 2018), a metabolic intermediates of one bacteria may act as the starting material of other bacteria (Surkatti and El-Naas, 2018), different genetic makeup (Woyke et al., 2006), synergetic effects of different microbial species, (Ghazali et al., 2004) or by synthesising different variant of catalytic enzymes. Bacterial cells in the microbial consortium can also combine their metabolic capabilities to utilize the common complex nutrient (Gilbert et al., 2003). Biodegradation of complex hydrocarbon mixture by the microbial consortium offers a combination of diverse enzymes which promotes the biodegradation processes (Wongwilaiwalin et al., 2010). However, microbial consortium immobilization for biodegradation of PAHs was not studied previously. The aim of the present study is to evaluate the naphthalene biodegradation performance by *Pseudomonas putida* strain KD10 and *Pseudomonas* sp. consortium as a free and immobilized format. Previous study showed that *Pseudomonas putida* strain KD9 cells decrease in size and shape from rod to sphere during naphthalene biodegradation and their specific growth rate was also little slower (Dutta et al., 2018). Keeping this fact in the mind, both morphological types of bacteria were applied in this present set of study. Additionally, the naphthalene 1, 2-dioxygenase (*nah*Ac) was sequenced and analyzed.

## 2. Materials and methods

### 2.1. Chemicals

Naphthalene was purchased from Sigma-Aldrich chemicals Pvt. Ltd. (USA) and all other chemicals used for media preparation were procured from HiMedia Laboratory (Mumbai, India). GC-MS and HPLC grade solvents were procured from Fisher Scientific (Mumbai, India). Sodium alginate (CAS No. 9005-38-3) of medium viscosity was purchased from Merck Pvt. Ltd. (USA).

### 2.2. Microorganisms, growth media, growth condition and consortium preparation

Soil samples were collected from petroleum refinery waste sites near Indian Oil (Haldia, West Bengal, India). The enrichment isolation and strain identification were carried out according to the standard protocol described previously (Dutta et al., 2017). Carbon deficient minimum medium (CSM), with a pH of 7.1, was used to cultivate the bacteria. Naphthalene was used as a sole source of carbon and energy in CSM with following compositions: 0.2 g L^-1^ MgSO_4_, 7H_2_O; 0.08 g L^-1^Ca(NO_3_)_2_, 4H_2_O; 0.005 g L^-1^ FeSO_4_, 7H_2_O; 4.8 g L^-1^, K_2_HPO_4_; 1.2 g L^-1^ KH_2_PO_4_. *Pseudomonas putida* strain KD6 (KX786159.1) and *Pseudomonas putida* strain KD9 (KX786158.1) and the newly isolated *Pseudomonas putida* strain KD10 (KX786157.1) were used to prepare the blend of *Pseudomonas* sp. consortium. The consortium was maintained in Luria-Bertani broth at 31°C with 150 rpm in order to grow the bacterial cell in its normal size and shape. Alternatively, bacterial cells were grown in CSM with naphthalene a as sole source of carbon and energy to obtain altered morphological variant.

### 2.3. Gene sequence analysis

#### 2.3.1. Detection of naphthalene 1, 2-dioxygenase and catechol 2, 3-dioxygenase

The conventional polymerase chain reaction (PCR) for naphthalene 1, 2-dioxygenase (*nah*Ac) and catechol 2, 3-dioxygenase (*nah*H) using specific primers (Table S1) were performed. Additionally, PCR product of *nah*Ac was sequenced and analyzed according to the methods described previously (Dutta et al., 2017).

#### 2.3.2. Clustering and phylogenetic analysis

The evolutionary distance of naphthalene 1, 2-dioxygenase among different bacterial species was analyzed using BLOSUM weighted matrix followed by pairwise distance computation using MEGA (v7.0) (Kumar et al., 2016). The distance matrix was then clustered using R (Team, 2013) to create the cladogram and heatmap. The phylogenetic position of the isolated *Pseudomonas putida* strain KD10 was analyzed using the previous method (Dutta et al., 2017).

#### 2.3.3. Molecular docking

The rigid-body molecular docking was conducted using Auto Dock (v4.2.1) (Morris et al., 1998). Briefly, the center grid dimensions were set to 20.271×61.989×87.168 with a grid spacing of 0.375 Å. The virtual screening was repeated for 10 times with the same unaltered docking parameters having 2.0 Å cluster tolerance. Additionally, the rigid-flexible molecular docking was performed using Molecular Operation Environment (Chemical Computing Group, Montreal Inc., Canada). The latter scoring function was employed to identify the most favourable docked poses and to estimate the binding affinity of the protein-ligand complexes. The non-covalent interactions were analyzed using the previous method (Salentin, S., et al., 2015).

### 2.4. Enzyme kinetic assay

The enzyme kinetic parameters of the naphthalene 1, 2-dioxygenase _I250, V256_ was performed using the cell-free extract of the *Pseudomonas putida* strain KD10, grown in 250 mL of CSM with naphthalene (500 mg L^-1^) as a sole source of carbon and energy as described by previously (Dutta et al., 2017).

### 2.4. Detection of solvent efflux pumps system

The detection of solvent efflux pump system (*srp*ABC) in *Pseudomonas putida* strain KD10 was performed using conventional polymerase chain reaction in a thermal cycler (Mastercycler® nexus gradient, Eppendrof, (Germany). Standard reaction mixtures were prepared using forward, reverse primers (Table S1) as described previously (Dutta et al., 2018).

### 2.5. Cell immobilization

#### 2.5.1. Cell immobilization in calcium alginate beads

*Pseudomonas putida* strain KD10 and *Pseudomonas* sp. consortium were immobilized according to the standard protocol described previously (Daâssi et al., 2014). Briefly as sodium alginate was dissolved in 0.9 wt. % NaCl (1 gm in 40 mL 0.9 wt. % NaCl) for 24 h and sterilized by autoclaving (121°C for 15 min). Two grams of bacterial cell mass was added to 8 ml of NaCl solution and again added to the sterile alginate solution. The mixture was then gently vortex for complete homogenization and extruded dropwise through a hypodermic syringe into chilled sterile CaCl_2_ solution (Figure 2). The beads were hardened in the same solution at room temperature with gentle stirring for 1 h. Finally, the beads were washed several times with 0.9 wt. % NaCl to remove excess calcium ions and free cells. The beads have an average diameter of 0.5 mm and stored at 4°C. Sterile beads (without microorganisms) were used to monitor the abiotic loss of naphthalene. Sodium alginate of medium viscosity (≥2,000 cP) was used to prepare the calcium alginate beads (CABs).

**Figure 1.**
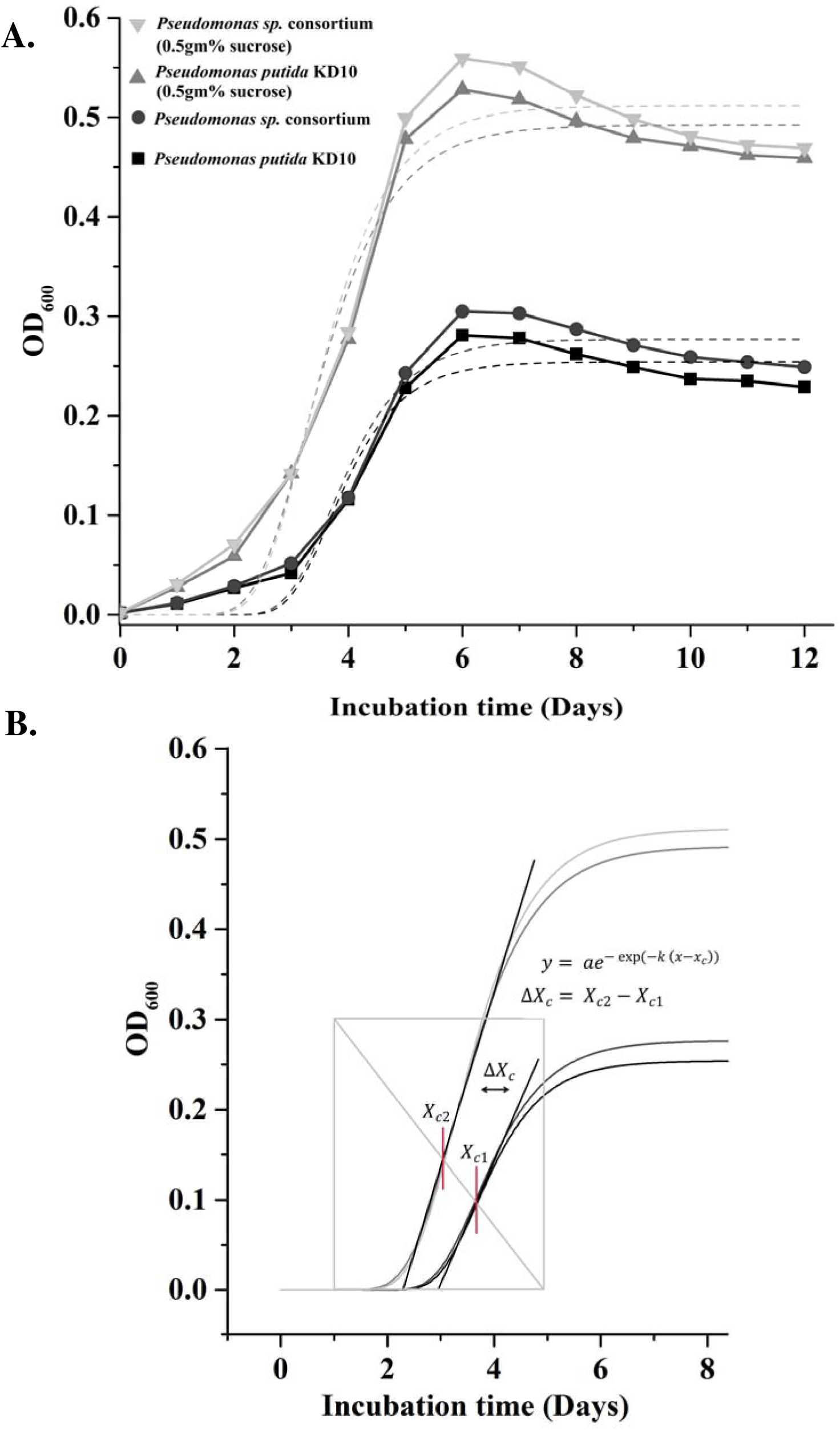
Growth kinetic model of *Pseudomonas putida* and *Pseudomonas* sp. consortium during naphthalene biodegradation. **A.** Gompertz’s growth kinetic model fit of the biodegradation of naphthalene by *Pseudomonas putida* strain KD10 and by the *Pseudomonas* sp. consortium. **B.** Effect of 0.5 gm. % sucrose supplementation on growth pattern of *Pseudomonas putida* strain KD10 and *Pseudomonas* sp. consortium.

**Figure 2.**
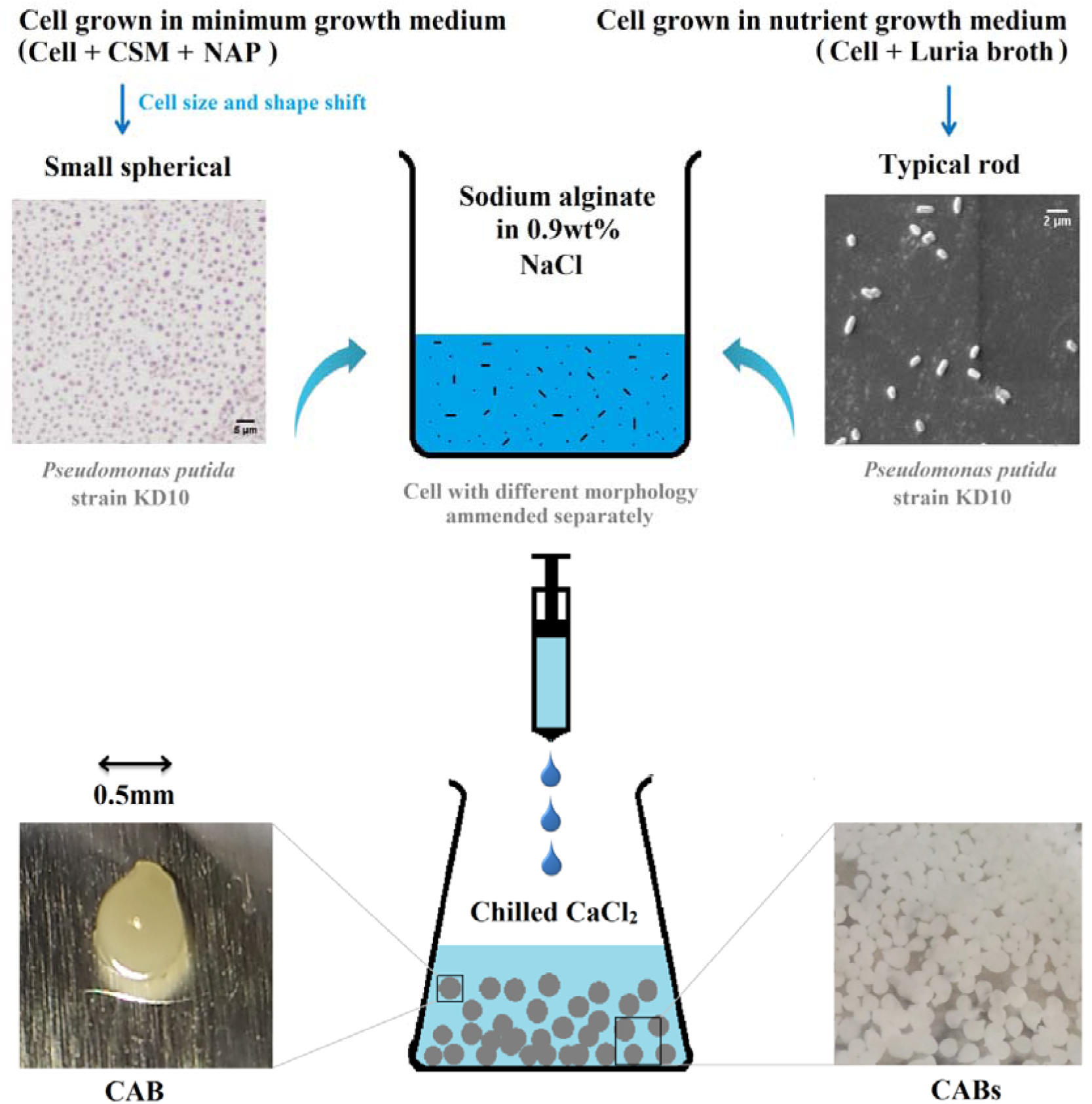
Cell immobilization in calcium alginate beads (CABs) with different cell morphology of the *Pseudomonas putida* strain KD10.

#### 2.5.2. Cell viability count

The viable cell enumeration in the CABs was performed by using a protocol described previously (Usha et al., 2010). In brief, CABs was washed in saline and keep submerged for 10 min (for saline soaking). Following soaking, the CABs were shaken with glass beads for 15 min and 1 g CABs were homogenized in 9 ml saline. The saline was then used for viable cell enumeration onto a nutrient agar plate. Plating was done with this treated saline by series dilution method up to 10^-5^ dilutions on nutrient agar plates. Plating was also done for the initial CFU count. For every dilution, 10 μl of the solution was plated. Plating was done by the pour plate method. Plates were incubated at 37°C for 24 h. (Jain and Pradeep, 2005).

### 2.6. Scanning electron microscopic (SEM) study

Samples for SEM analysis were prepared by a protocol described elsewhere (Dutta et al., 2018). The dried samples were coated in a sputter coater (Quorum-SC7620) under vacuum with a thin gold layer right before SEM analysis using a scanning electron microscope (Zeiss, EVO 18, Germany) with an accelerating voltage of 5 kV.

### 2.7. Biodegradation kinetics

#### 2.7.1. Biodegradation of naphthalene in the liquid medium

Naphthalene biodegradation study was conducted by the protocol described previously (Dutta et al., 2017). The conical flasks were incubated at 31°C with 150 rpm and uninoculated conical flask were used as control. Culture medium was collected at regular interval of 72 h for degradation and growth kinetic study. Additionally, the effect of initial concentration of naphthalene (150-2500 mg L^-1^) was studied with different immobilized systems.

#### 2.7.2. Chemical analysis

Naphthalene biodegradation was analyzed by using the 1260 infinity series HPLC system (Agilent, Santa Clara, CA, USA) equipped with Zorbax SB-C18 reversed-phase column (4.6 × 12.5 mm, 5 µm). The analysis of naphthalene concentrations was conducted using isocratic elution conditions with the mobile phase 80:20 (v/v) methanol: water at a flow rate of 1 ml min^-1^. The detection was performed at 254 nm according to the protocol described in the literature (Dutta et al., 2017). Additionally, the metabolic intermediates of naphthalene biodegradation were analyzed using the GC-MS system (GC Trace GC Ultra, MS-Polarisq, Thermo Scientific India Pvt. Ltd) equipped with a capillary column (TR-WaxMS, 30 m × 0.25 mm [ID] × 0.25 µm film thickness) by the protocol described elsewhere (Dutta et al., 2017). The entire analysis was performed in electron ionization, at full scan mode. The metabolite identification was based on the mass spectra comparison using the NIST Mass Spectral library (v2.0, 2008).

### 2.8. Data analysis

The first-order degradation kinetics model was used to estimate the residual naphthalene in CSM using equations 1, the algorithms as expressed determine the half-life (*t*_1/2_) values of naphthalene in CSM. The substrate inhibition kinetic parameters were calculated using equation 2.

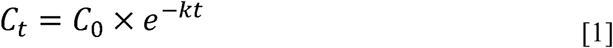

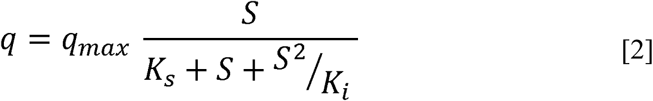

The effect of SCS on growth pattern was measured by calculating the difference between optical densities at 600_nm_ and expressed by slight extension (Equation 4) of the Gomperz’s sigmoid growth fit equation (Equation 3).

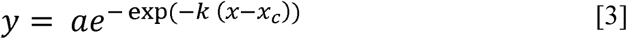

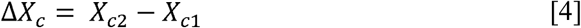

The growth pattern change by SCS was considered as CFU shift (Equation 5). Where, X_c2_ = Optical density at X_c2_ and X_c1_ = Optical density at X_c1_.

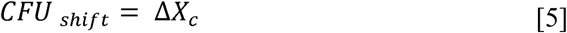

The bacterial growth kinetics were analyzed by applying the Gomperz’s model using Levenberg-Marquardt algorithm in Origin® 2016 (California, USA) and the degradation kinetics were analyzed by using Graphpad Prism® 6.01 (San Diego, CA, USA).

## 3. Results and discussion

### 3.1. Isolation, identification and growth patterns of NAP degrading bacterial strain

Pseudomonas putida strain KD10 was isolated from petroleum refinery waste with its distinguished colony morphology on CSM agar plate (Figure S1a, b). The PCR product of 16S rRNA gene of the strain KD10 was sequenced. Moreover, the multiple sequence alignment followed by phylogenetic assessment suggests that the strain belongs to the *Pseudomonas putida* group and it showed 99% sequence similarity with the previously deposited sequences of *Pseudomonas putida* (Figure S2). The 16S rRNA gene sequence was deposited at NCBI (https://www.ncbi.nlm.nih.gov/) under the GenBank accession number KX786157.1.

Naphthalene biodegradation by the *Pseudomonas putida* KD10 was preliminarily confirmed by the catechol test (Figure S1C) followed by growth pattern analysis in CSM with naphthalene as a sole source of carbon and energy. Additionally, growth pattern with 0.5 gm % sucrose as a secondary carbon supplement (SCS) was compared with growth pattern of non-supplemented CSM. Gompertz’s model fit (Equation 3) was used to analyze the growth curve (Table S4).

### 3.2. Pseudomonas sp. consortium, growth pattern and NAP biodegradation

The growth pattern of the *Pseudomonas* sp. consortium, as shown in Figure 1 was optimized at 31°C and pH 7.1 which reveal successful cooperation among *Pseudomonas putida* strain KD6, KD9 and KD10. Further, the growth curve model indicates a significant increase in total biomass of the *Pseudomonas* sp. consortium and they also showed a high concentration of naphthalene tolerance potential.

### 3.3. Gene sequence and phylogenetic analysis

Naphthalene 1, 2-dioxygenase (*nah*Ac), encoded by *Pseudomonas putida* strain KD10 have only two point mutations at I250, and V256 which were replaced by methionine and glycine respectively (Figure 3A, S6). However, additional mutations were found in the same chain A at K200, A210, E264, M284 and N334 by replacing glutamic acid, glycine, aspartic acid, isoleucine, respectively (Dutta et al. 2017, 2018). The three dimensional structural analysis suggests that the mutations at 200, 210, 284 and 334 causes a little structural change in comparison to the wild type variant of naphthalene 1, 2-dioxygenase (Figure 3B). Conversely, the amino acids stretch at the close proximity of the active site residues exactly from 248 to 266 showed a structural mismatch among all three mutant and wild type variants of naphthalene 1, 2-dioxygenase (Figure 3B). The active site of an enzyme tends to evolve fast to attain its maximum performance and functionality in a particular environment (James and Tawfik, 2003).

**Figure 3.**
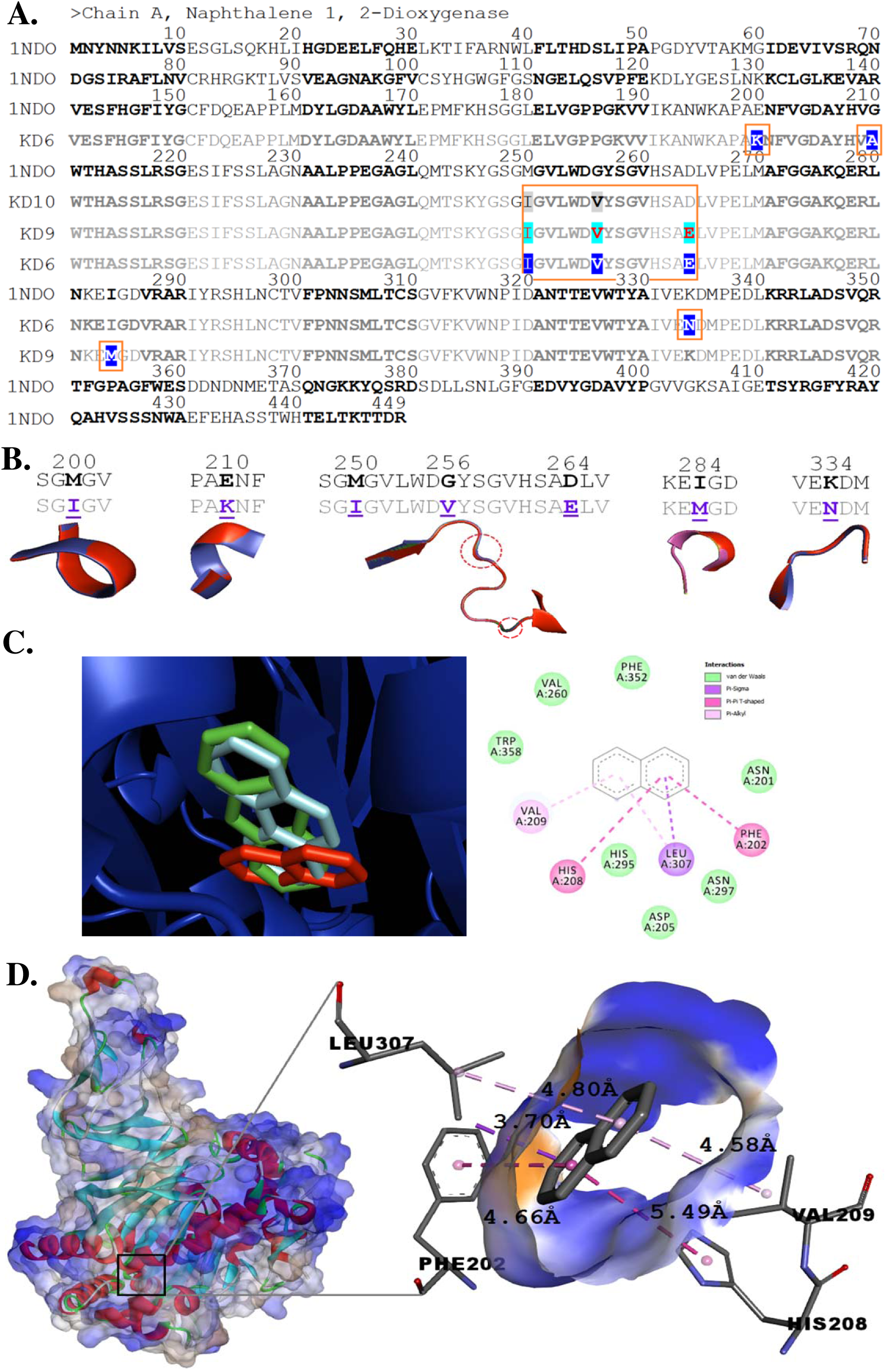
Sequence and structure analysis of the naphthalene 1, 2-dioxygenase (*nah*Ac) encoded by *Pseudomonas putida*. **A**. The amino acid sequence of wild type and three mutant variants of the naphthalene 1, 2-dioxygenase (*nah*Ac). Mutated residues are highlighted in blue and the common mutation prone amino acid stretch is highlighted with red box. **B.** Structural mismatches among mutant variants and wild type naphthalene 1, 2-dioxygenase. Each mutated residues and its subsequent neighbour residues of all the mutant variant of naphthalene 1, 2-dioxygenase is overlaid with cartoon representation. The highlighted red circle indicates local mismatch among the mutant variant of naphthalene 1, 2-dioxygenase **C.** Different ligand binding posture of the naphthalene 1, 2-dioxygenase encoded by *Pseudomonas putida* strain KD10. Naphthalene (red), phenanthrene (cyan) and anthracene (green) binding postures with major interacting amino acid residues (left). Two-dimensional presentation of the interacting residues of *nah*Ac_I250, V256_ (right). **D.** Docking pose of naphthalene 1, 2-dioxygenase (Chain A) encoded by *Pseudomonas putida* strain KD10. The four major naphthalene interacting residues and their molecular distance are labelled in black.

The alteration of amino acids bases from 248 to 266 of chain A, was a common feature in all three mutant variants of *nah*Ac and this particular stretch of amino acid is very close to the active site residues (Figure 3C). This implies that the “248-266” amino acid stretch is highly prone to mutation and it may influence on the enzymatic efficiency and environmental adaptability. Thus, our study provides a new insight, which could be beneficial for rational approaches of enzyme redesigning.

The biodegradation performance of a bacterial strain is linked with the three dimensional structure of the catalytic enzyme and ligand binding posture (Singh et al., 2019). Moreover, alteration of a single amino acid in the catalytic domain of the enzyme caused different ligand binding postures which eventually lead to a unique biodegradation pathways (Ferraro et al., 2006). In addition, site directed mutagenesis in the catalytic domain offers superior enzyme activity (Parales et al., 1999; Parales, 2003). The comparative analysis of binding free energy helps to comprehend the superior naphthalene biodegradation performance by the *Pseudomonas* sp. consortium, which is, in fact, the summative activity of all three mutant variants of naphthalene 1, 2-dioxygenase.

The evolutionary trace on the naphthalene 1, 2-dioxygenase among different species were studied through pairwise distance matrix analysis (Figure 4), which suggests a significant intra and inter species difference among *Sphingopyxis sp.*, *Croceicoccus naphthovorans, Burkholderia multivorans, Burkholderia sp. Massilis sp. and Cycloclasticus sp.* Conversely, other *Pseudomonas* sp. particularly, *Pseudomonas benzenivorans, Pseudomonas balearica, Pseudomonas stutzeri, Pseudomonas kunmingensis, Pseudomonas frederikbergensis* have similarities in their version of naphthalene 1, 2-dioxygenase. A few other species such as *Xenophilusa zovorans, Ralsoniam annitolilytica*, and *Paraburkholderia aromaticivorans* does not show significant evolutionary distance in their version of naphthalene 1, 2-dioxygenase (Figure 4). However, a few strains of *Burkholderria multivorans* showed similarities and some other does not (Figure 4). The source of collection of these two *Burkholderria multivorans* species may be the reason for such variation (Li et al., 2007). In a study by (Su et al., 2016), the epigenetic impacts on the metabolic enzymes have been suggested. Moreover, the divergence of evolutionary distances of the same enzyme between different and same bacterial species indicates thrive in a particular microenvironment. However, further studies are needed to identify the underlying cause and mechanisms of such variation intra-species variation.

**Figure 4.**
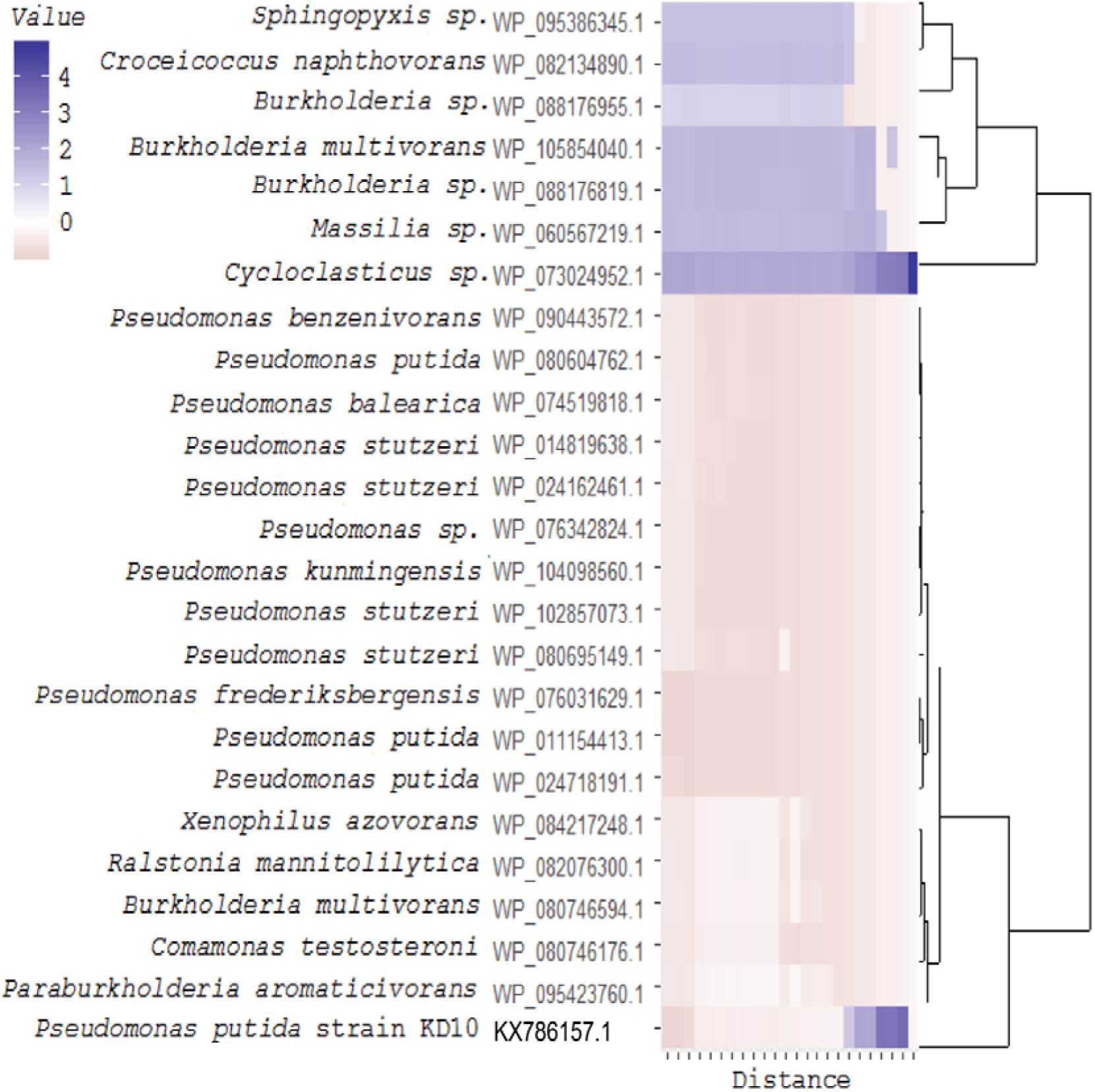
Distance matrix and cladogram of the naphthalene 1, 2-dioxygenase. The evolutionary distance of the naphthalene 1, 2-dioxygenase I_250_, V_255_ variant encoded by *Pseudomonas putida* strain KD10 other variants by different microbial species is scaled by colour index (upper-left side).

### 3.4. Molecular docking

The result depicted from the rigid body molecular docking, suggests *nah*Ac-KD10_I250, V256_ mutant has a little higher binding free energy than that of mutant *nah*Ac encoded by KD9 but little lower than the *nah*Ac mutant variant encoded by the KD6 (Table 1). However, the rigid-flexible molecular docking using MOE algorithms showed *nah*Ac I_250_, G_256_ mutant variant has ΔG_London_ of −7.1 kcal mol^-1^ and ΔG_GBVI/WSA_ of −1.68 kcal mol^-1^, which is little higher than other two mutant variants of *nah*Ac encoded by KD6 and KD9 (Table 1). The interacting amino acid residues in the active site were confined to be quite same in three mutant variants, except one variation, *i.e.*, His_208_ which is located about 5.49 Å away from the bicyclic ring of naphthalene (Figure 3D). The altered binding free energy of the *nah*Ac-KD10_I250, V256_ may be due to the fact that the two altered amino acid residues reside in “248-266” and participation of His_208_ as a unique interacting residue (Figure 3C). Furthermore, the non-covalent interactions of the *nah*Ac-KD10_I250, V256_ with naphthalene, phenanthrene and anthracene suggest that Phn 352 was common residue and His 208 is involve in π- staking with a bond angle of 80.58° and 77.19° respectively for naphthalene and anthracene (Table S5).

**Table 1.**
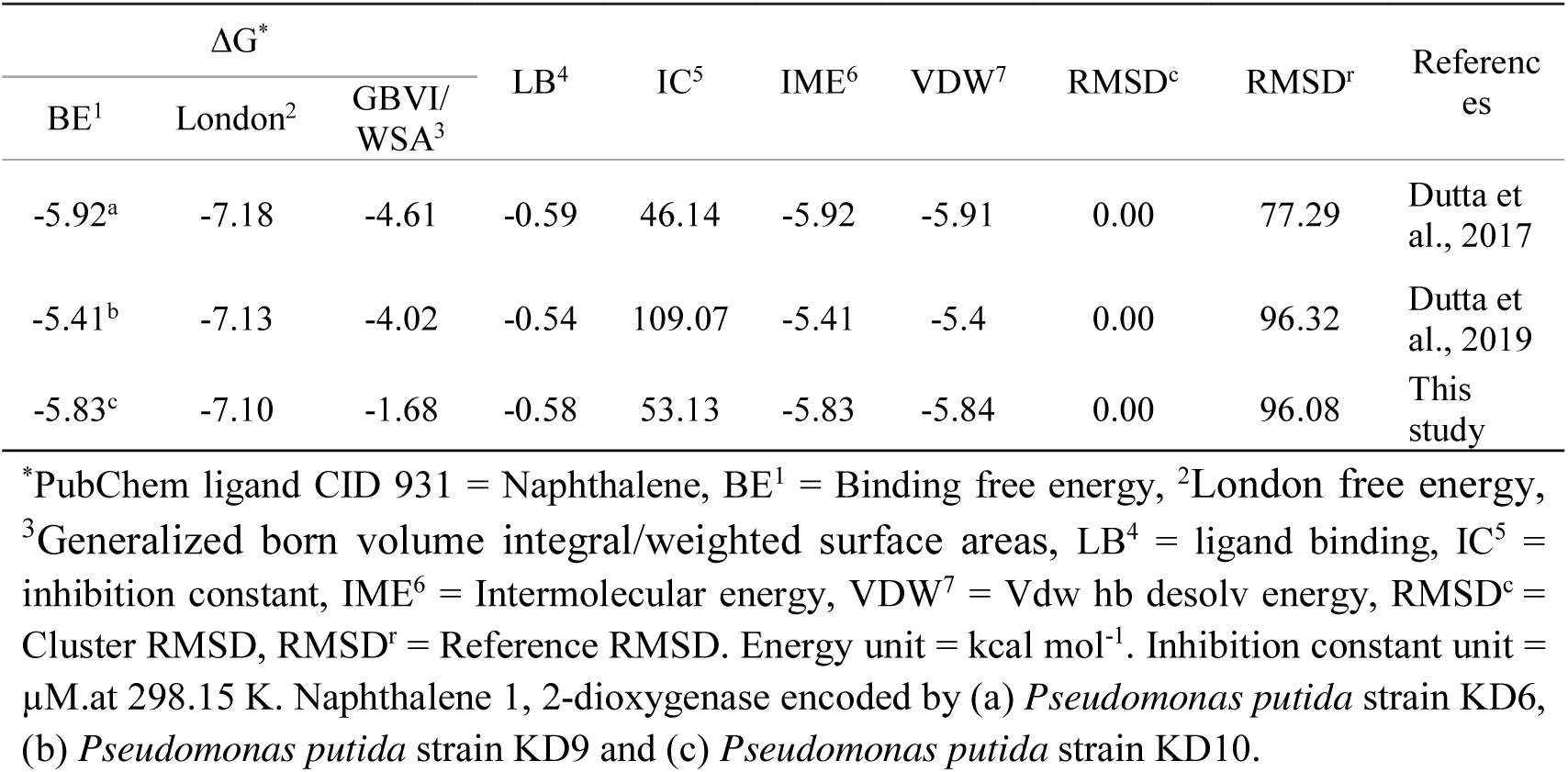
Summary of rigid-flexible molecular docking for the analysed mutant naphthalene 1, 2-dioxygenase encoded by different *Pseudomonas putida* strains.

| $\Delta G^*$ | | | LB<sup>4</sup> | IC<sup>5</sup> | IME<sup>6</sup> | VDW<sup>7</sup> | RMSD<sup>c</sup> | RMSD<sup>r</sup> | References |
| --- | --- | --- | --- | --- | --- | --- | --- | --- | --- |
| BE<sup>1</sup> | London<sup>2</sup> | GBVI/WSA<sup>3</sup> |  |  |  |  |  |  |  |
| -5.92<sup>a</sup> | -7.18 | -4.61 | -0.59 | 46.14 | -5.92 | -5.91 | 0.00 | 77.29 | Dutta et al., 2017 |
| -5.41<sup>b</sup> | -7.13 | -4.02 | -0.54 | 109.07 | -5.41 | -5.4 | 0.00 | 96.32 | Dutta et al., 2019 |
| -5.83<sup>c</sup> | -7.10 | -1.68 | -0.58 | 53.13 | -5.83 | -5.84 | 0.00 | 96.08 | This study |

\*PubChem ligand CID 931 = Naphthalene, BE<sup>1</sup> = Binding free energy, <sup>2</sup>London free energy, <sup>3</sup>Generalized born volume integral/weighted surface areas, LB<sup>4</sup> = ligand binding, IC<sup>5</sup> = inhibition constant, IME<sup>6</sup> = Intermolecular energy, VDW<sup>7</sup> = Vdw hb desolv energy, RMSD<sup>c</sup> = Cluster RMSD, RMSD<sup>r</sup> = Reference RMSD. Energy unit = kcal mol<sup>-1</sup>. Inhibition constant unit = $\mu$ M.at 298.15 K. Naphthalene 1, 2-dioxygenase encoded by (a) *Pseudomonas putida* strain KD6, (b) *Pseudomonas putida* strain KD9 and (c) *Pseudomonas putida* strain KD10.

### 3.5. Enzyme kinetic assay

Enzyme activity by cell free extract of Pseudomonas putida KD10 was studied using Naphthalene as substrate. The cell free extract of strain KD10 degraded Naphthalene, with *R*^2^ of 0.935 indicating the experimental data are well correlated with the model (Table 2). The performance constant (Kcat/ Km) was 0.142×10^3^ ml^-1^ mol.^-1^ s^-1^ which further validate the chemical data obtained from chemical analysis (Table 1). In previous study, the performance constant of *Pseudomonas putida* KD6 was little higher than *Pseudomonas putida* strain KD10, suggesting the six altered amino acid residues of the mutant naphthalene 1,2-dioxygenase of the KD6 may be main cause of this difference (Dutta et al. 2017). The performance constant (Kcat/ Km) is the indicator of the performance of an enzyme and usually the higher Kcat/Km meaning better enzymatic performance (Koshland Jr DE, 2002).

**Table 2.**
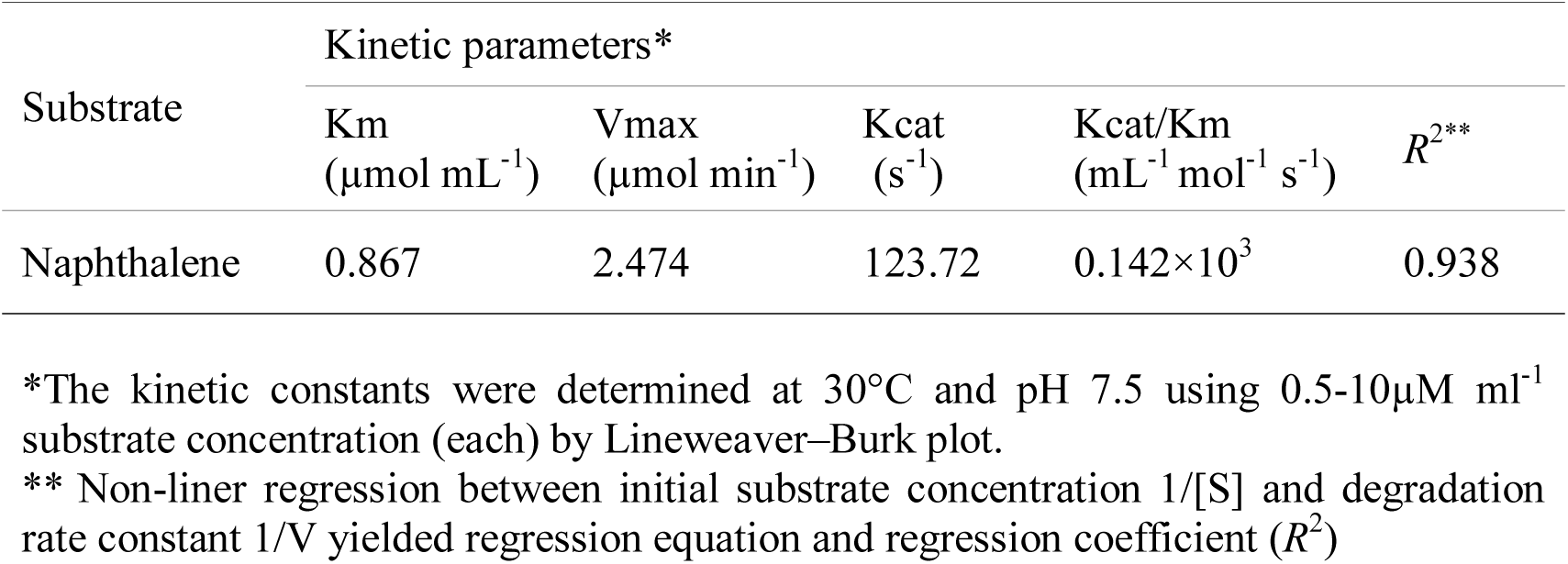
Apparent enzyme kinetic parameters of naphthalene 1, 2-dioxygenase _I250, V256_ encoded by *Pseudomonas putida* strain KD10.

| Substrate | Kinetic parameters* |  |  |  |  |
| --- | --- | --- | --- | --- | --- |
| | K<sub>m</sub><br>($\mu\text{mol mL}^{-1}$) | V<sub>max</sub><br>($\mu\text{mol min}^{-1}$) | K<sub>cat</sub><br>($\text{s}^{-1}$) | K<sub>cat</sub>/K<sub>m</sub><br>($\text{mL}^{-1} \text{mol}^{-1} \text{s}^{-1}$) | $R^{2**}$ |
| Naphthalene | 0.867 | 2.474 | 123.72 | $0.142 \times 10^3$ | 0.938 |

\*The kinetic constants were determined at 30°C and pH 7.5 using 0.5-10 $\mu\text{M mL}^{-1}$ substrate concentration (each) by Lineweaver–Burk plot.
\*\* Non-linear regression between initial substrate concentration 1/[S] and degradation rate constant 1/V yielded regression equation and regression coefficient ( $R^2$ )

### 3.6. Detection of solvent efflux system

The cellular microenvironment and internal homeostasis are crucial for maintaining normal cellular functions and cell tends to adopt several strategies in order to achieve it (Blanco et al., 2016). One such adaptation strategy is the solvent efflux pump system, that control intracellular toxicity to some extent (Kusumawardhani et al., 2018). The role of *srp*ABC solvent efflux pump system in the biodegradation of PAHs are common in literature (Bugg et al., 2000). Besides the other cellular activities, *srp*ABC also assist in gaining antimicrobial resistance (Schweizer, 2003). Biodegradation studies on chlorpyrifos indicate that the intermediates formed during biodegradation, act as an antimicrobial agent to other species, meaning that the metabolic intermediates may have a role on inter-species competition in a particular micro-environment to thrive in nutrient-limiting condition (Anwar et al., 2009), (Raes and Bork, 2008). Besides, the role of *srp*ABC on biodegradation enhancement process is still poorly understood. The presence of *srp*ABC in *Pseudomonas putida* strain KD10 (Figure S5) and the efficient naphthalene biodegradation property can be interlinked (Dutta et al., 2018).

### 3.7. Cell immobilization and viability count

The viscosity of CABs determines its efficiency of the immobilization and its performance of detoxification of environmental pollutant (Young et al., 2006). Therefore, sodium alginates of medium viscosity were chosen for this study. The efficiency of cell immobilization in CABs was evaluated by enumeration of viable cells (Table S3). Bacteria immobilized in CABs do not significantly lose its viable biomass after 21 days of incubation at 4°C.

### 3.8. Naphthalene biodegradation studies

#### 3.8.1 By free Pseudomonas putida strain KD10

Biodegradation kinetics of naphthalene (500 mg L^-1^) by the *Pseudomonas putida* strain KD10 was first tested to determine its naphthalene biodegradation potential (Figure 4). It was found that after 12 days of incubation at 31°C the amount of residual naphthalene reduced significantly (*p*<0.05) (Figure 5). Results are also summarised at Table 3 along with names of the immobilized systems. The first order degradation kinetic data suggest that at the end of the incubation *Pseudomonas putida* strain KD10 efficiently decompose 95.22 % naphthalene in CSM as a sole source of carbon and energy. The values of degradation rate constant (*k*) and half-life (t_1/2_) were 0.2193 and 3.1 days with *R*^2^ of 0.981. In the previous study, 99.1 % of initial naphthalene was removed within 96 h by strain *Bacillus fusiformis* (BFN). However, the initial concentration was very low (50 mg L^-1^ of initial naphthalene) (Lin et al., 2010).

**Figure 5.**
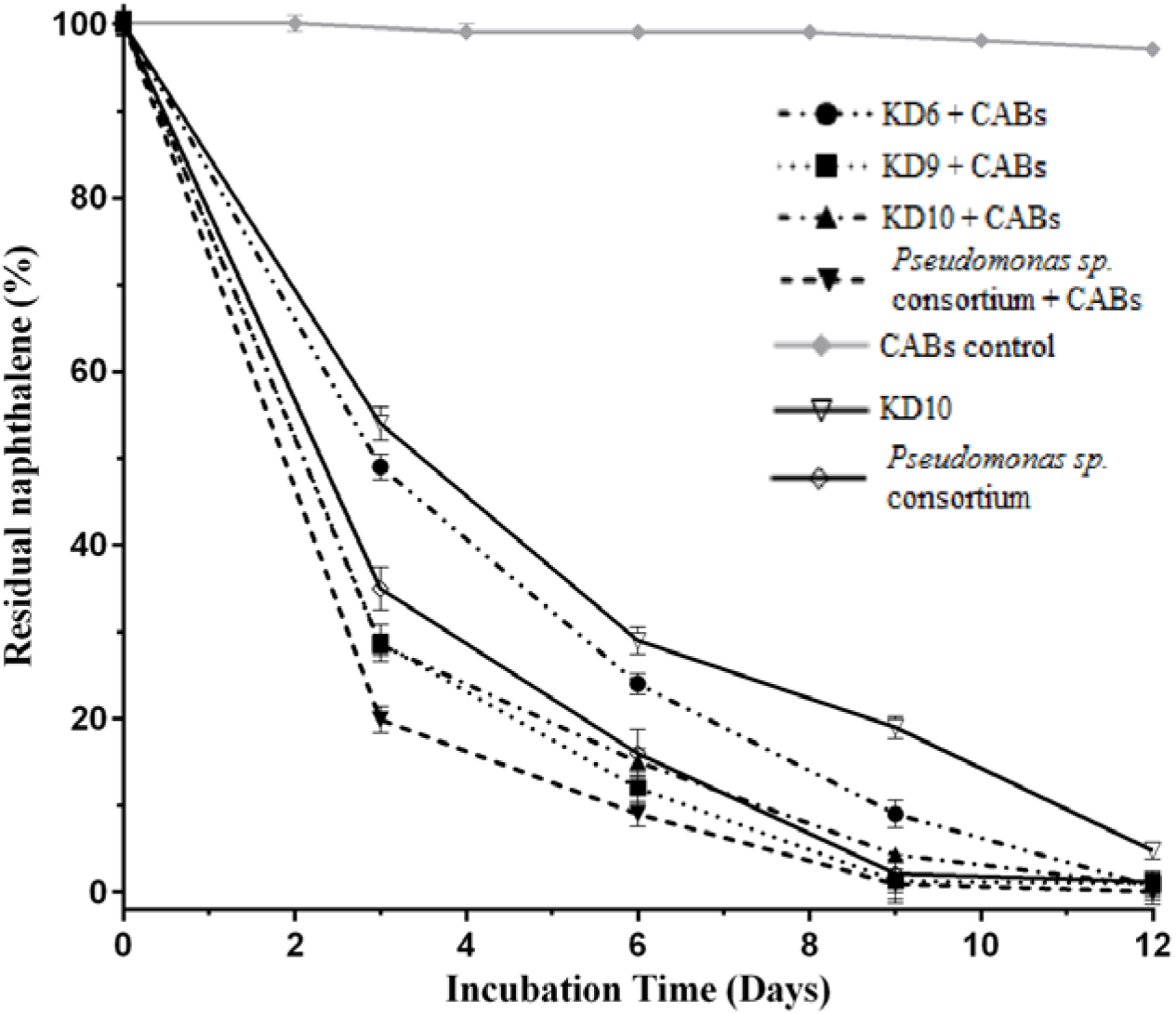
Naphthalene degradation kinetics of *Pseudomonas putida* strain KD10 and *Pseudomonas* sp. consortium free and cell immobilized in calcium alginate beads.

**Table 3.**
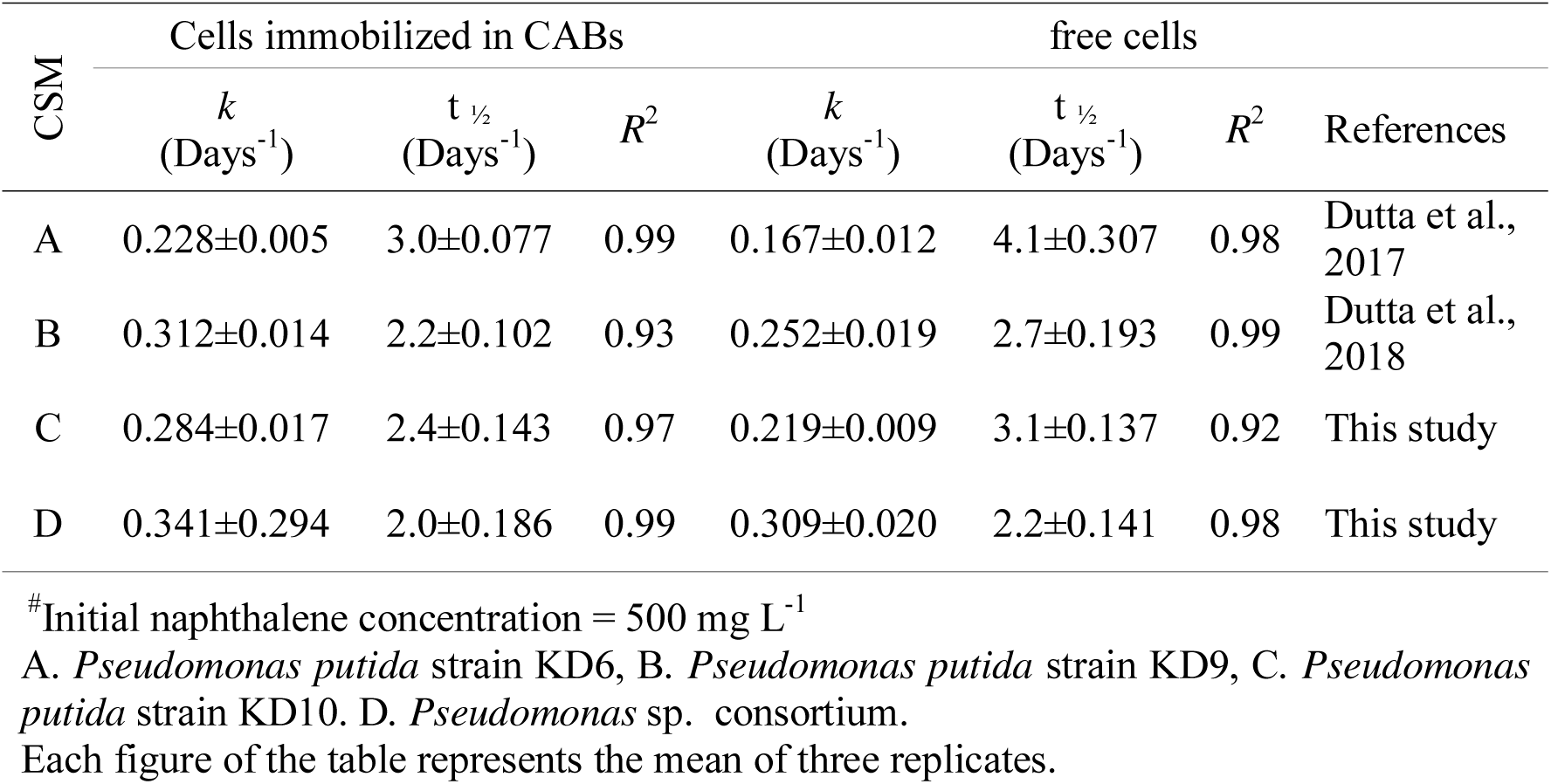
Parameters of first-order biodegradation kinetics of naphthalene by different free and immobilized *Pseudomonas putida* strains.

| CSM | Cells immobilized in CABs |  |  | free cells |  |  | References |
| --- | --- | --- | --- | --- | --- | --- | --- |
| | $k$<br>(Days<sup>-1</sup>) | $t_{1/2}$<br>(Days<sup>-1</sup>) | $R^2$ | $k$<br>(Days<sup>-1</sup>) | $t_{1/2}$<br>(Days<sup>-1</sup>) | $R^2$ | |
| A | 0.228±0.005 | 3.0±0.077 | 0.99 | 0.167±0.012 | 4.1±0.307 | 0.98 | Dutta et al., 2017 |
| B | 0.312±0.014 | 2.2±0.102 | 0.93 | 0.252±0.019 | 2.7±0.193 | 0.99 | Dutta et al., 2018 |
| C | 0.284±0.017 | 2.4±0.143 | 0.97 | 0.219±0.009 | 3.1±0.137 | 0.92 | This study |
| D | 0.341±0.294 | 2.0±0.186 | 0.99 | 0.309±0.020 | 2.2±0.141 | 0.98 | This study |

<sup>#</sup>Initial naphthalene concentration = 500 mg L<sup>-1</sup>
A. *Pseudomonas putida* strain KD6, B. *Pseudomonas putida* strain KD9, C. *Pseudomonas putida* strain KD10. D. *Pseudomonas* sp. consortium.
Each figure of the table represents the mean of three replicates.

After confirmation of naphthalene biodegradation potential of *Pseudomonas putida* strain KD10, the stain was allowed to grow in association with other two strains of *Pseudomonas putida*, namely *Pseudomonas putida* strain KD6 and *Pseudomonas putida* strain KD9 (Dutta et al., 2017; Dutta et al., 2018). These two strains were selected to develop the blend of *Pseudomonas* sp. consortium, because they were collected from different isolation points, they encode different variant of *nah*Ac and they have common optimized growth parameters (temperature 31°C and pH 7.1) (Dutta et al., 2017 & Dutta et al., 2018). Further, *Pseudomonas putida* strain KD6 encodes a six point mutant *nah*Ac with the capability to co-degrade high concentration of naphthalene, phenanthrene (PHN) and pyrene (PYR), (500 mg L^-1^ each). Moreover, the stain KD9 encodes a four point mutant variant of *nah*Ac with rhamnolipid production capabilities (Dutta et al. 2018).

In the previous study, *t*_1/2_ of naphthalene co-biodegradation with phenanthrene and pyrene by *Pseudomonas putida* strain KD6 was 4.1 days, which was significantly reduced to 2.2 days when bacterial cells were allowed to grow as *Pseudomonas* sp. consortium with only naphthalene (Table 3). Further, this value was found as 2.7 days for *Pseudomonas putida* KD9, suggesting *Pseudomonas putida* strain KD6 might face some sort of substrate inhibition by the co-presence PHN and PYR in the system (Jiang et al., 2018) and the cooperative nature of the *Pseudomonas* sp. consortium helps to enhance the naphthalene bio-utilization. The microbial consortium works several ways, *viz.*, by the division of labor, cross-feeding, *etc.* (Smid and Lacroix, 2013). Moreover, a successful consortium could also overcome shortcomings of single bacteria (Bhatia et al., 2018). However, it is a fact that, bacteria select PAH among mixed PAHs based on their structural simplicity first (Dutta et al., 2017) and the velocity could be optimized by reducing the initial concentration of the PAHs gradually according to the structural simplicity (Jiang et al., 2018).

#### 3.8.2. By immobilized Pseudomonas sp. consortium

The biodegradation kinetic of naphthalene by individual *Pseudomonas putida* strain KD6, KD9, KD10 and *Pseudomonas* sp. consortium immobilized in CABs, depicted further enhancement of overall naphthalene bio-utilization (Table 3). Cell immobilization using hydrogel, such as CAB found to be advantageous rather than free cells (Hameed and Ismail, 2018).

This phenomenon may be attributable due to the increased level of tightness of the cross-linked polymers of the calcium alginate beads that render bacteria adequate amount of protection from harsh environment (Chen et al., 2013). However, the free bacterial cell lacks the capabilities to degrade a high initial concentration of the toxicant because they followed the conventional growth phases (Marrot et al., 2006). In addition, exposure of free bacterial cells to the high initial concentration of toxicants may challenge them to experience shock-concentration (Zhao et al., 2006). Conversely, cells immobilized in calcium alginate beads can tolerate high concentration of the toxicant and decrease the lag phase duration (Kao et al., 2014). Moreover, the diffusion limitation natures of the CABs matrix provide a high local concentration of the cell population (Bezbaruah et al., 2009). Furthermore, CABs provides remarkable stability and reusable features that effectively reduces the production cost (Daâssi et al., 2014).

Biodegradation of naphthalene by immobilized *Pseudomonas* sp. consortium significantly elevated the overall naphthalene bio-utilization efficiency with t_1/2_ and *R*^2^ values of 2 days and 0.998 respectively, suggesting experimental data are well correlated to the model (Table 3). The individual cell population also displayed improved biodegradation efficiency. However, the biodegradation efficiency was maximum by *Pseudomonas putida* strain KD6, which showed a marked reduction of *t_1/2_* from 4.1 to 3.0 days (Table 3). It promptly suggests that CABs facilitate KD6 optimized naphthalene bio-utilization, that might be masked-up by the co-presence of NAP, PHN, PYR (Jiang et al., 2018). However, further studies are required to investigate behavioural patterns of a consortium with mixed PAH.

Bacterial cell grown on CSM with naphthalene as a sole source of carbon and energy showed a delayed growth rate and their total biomass was also low (Figure 1). Nevertheless, growing cells on CSM prior to immobilization in CABs provide them essential adaptation to the toxicant and mature them for such stress condition (Table 3). Immobilized cell system was useful for bioremediation of a toxicant after prior adaptation to the surrounding environment (Partovinia and Rasekh, 2018). However, growing cells in Luria-Bertani broth does not adapt cell adequately, and their naphthalene removal performance was very poor (Figure S3). The inoculum was normalized (OD_600nm_ = 0.002) for each *Pseudomonas putida* strain in order to prepare the *Pseudomonas* sp. consortium. In aqueous medium, the immobilized bacterial cell mainly on the surface was exposed to naphthalene and primarily involved in the naphthalene bio-utilization process. However, due to the micro-porous feature of the calcium alginate beads, microbial cells immobilized other than surface are also participates in the bio-utilization and bio-sorption process of naphthalene. Further, with time due to the mechanical force generated by shaking, a few microbial cells may release from the calcium alginate beads. However, to fuel up the cell, it is necessary to uptake naphthalene for bio-utilization through step by step intracellular enzymatic reactions (Lin et al., 2014).

#### 3.8.3. Effect of initial naphthalene concentration on biodegradation kinetics

The effect of different initial concentration of naphthalene (150 – 2500 mg L^-1^) on degradation kinetics was evaluated with immobilized KD6, KD9, and KD10 as individual and as a consortium (Table S2). The results suggest cell immobilization in CABs facilitates bacteria to cope with a high initial concentration of NAP (Table S2). The substrate inhibition kinetic parameters, *viz.*, the maximum specific degradation rate (*q*_max_), and inhibition constant (*k*_i_) were highest for immobilized *Pseudomonas* sp. consortium and these values were 0.707 h^-1^ and 1475 mg L^-1^ respectively (Table S2). However, we did not find any significant change on half saturation constant (*k*_s_), suggesting the reaction does not depend on its initial concentration and the biodegradation process follows pseudo-first order reaction kinetics. In previous study, free *Pseudomonas putida* KD9 in CSM was capable to tolerate relatively low initial concentration of naphthalene with *k_i_* value of 1107 mg L^-1^ and addition of sucrose as SCS provides quite similar potential of naphthalene tolerance (*k_i_* of 1429 mg L^-1^) with that of immobilized *Pseudomonas* sp. consortium (Dutta et al., 2018). However, sucrose (0.5 gm. %) in the mobile open water system would not be beneficial for bacteria to overcome the high shock concentration of the toxicant, again suggesting cell immobilization and development of effective microbial consortium as systematic optimization of biodegradation process.

### 3.9. Detection of metabolic end products

The metabolic pathway of naphthalene biodegradation that might be followed by *Pseudomonas putida* strain KD10 was elucidated through GCMS analysis of the metabolites (Figure S4). In the previous study, the major metabolites ware restricted to be salicylaldehyde, catechol, D-gluconic acid and pyruvic acid (Dutta et al., 2018). However, in the present study, we have detected salicylic acid as an additional metabolite (Table 4). Catechol step into the TCA cycle by two possible pathway one is ortho-cleavage and another is meta-cleavage pathway. The meta-cleavage pathway is led by catechol-2, 3-dioxygenase (*nah*H), and presence of *nah*H (Figure S5) suggesting in *Pseudomonas putida* stain KD10 opt the meta-cleavage pathway. Furthermore, we assume that D-gluconic acid enters in the bio-conversion of naphthalene, possibly from glucose as a precursor (Figure 6) and it is finally converted to pyruvate via aldolase. D-gluconic acid implies impotence between bacteria-plant mutualistic association e.g., induction of phosphate solubilization processes (Rodriguez et al., 2004) which can inhibit fungal growth (Kaur et al., 2006). The metabolic intermediates *viz.*, salicylic acid, catechol has plant-growth promoting activity (Lee et al., 2010) e.g., the antioxidant property of the catechol help and promotes seed germination (Schweigert et al., 2001). The presence of d-gluconic acid as a major metabolite and its subsequent entry in TCA cycle from glucose precursor, suggest that KD10 may able to sequestered catechol upon needs to promote the plants’ growth (Figure 6).

**Figure 6.**
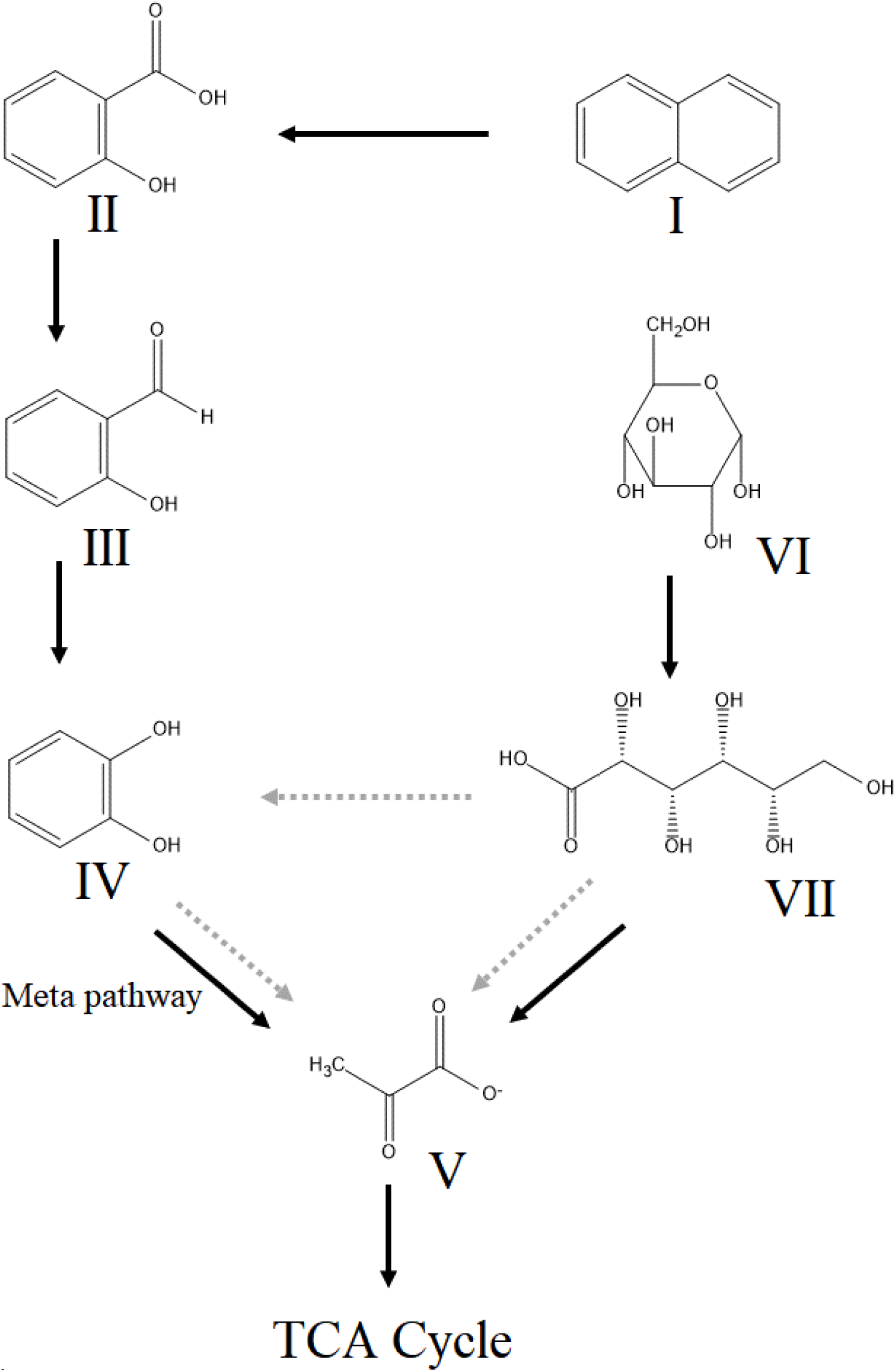
Proposed pathway of naphthalene biodegradation by *Pseudomonas putida* strain KD10. Naphthalene (I), salicylic acid (II), salicylaldehyde (III), catechol (IV), pyruvate (V), glucose (VI), d-gluconic acid (VII). The grey arrow indicates alternative pathway via d-gluconic acid.

**Table 4.** Major metabolite detected by GCMS analysis during naphthalene biodegradation by *Pseudomonas putida* KD10.

| Metabolite | R<sub>t</sub><br>(min) | Major ion peaks (m/z) | Suggested structure |
| --- | --- | --- | --- |
| I | 9.41 | 64.19, 92.17, 120.21, 138.22, 169.14, 191.18, 211.10 | Salicylic acid |
| II | 15.27 | 65.17, 74.41, 104.73, 121.10, 122.57, 142.21, 174.80, 201.22, 249.31 | Salicylaldehyde |
| III | 18.71 | 64.29, 81.35, 91.61, 110.47, 131.47, 149.71 | Catechol |
| IV | 31.89 | 61.21, 73.44, 76.24, 104.18, 117.41, 133.27, 177.23 | D-gluconic acid |
| V | 10.47 | 43.21, 61.27, 71.37 | Pyruvic acid |

R<sub>t</sub> = Retention time

## 4. Conclusion

Naphthalene biodegradation by free and immobilized *Pseudomonas putida* strain KD10 and *Pseudomonas* sp. consortium were studied. HPLC analysis showed 80.1% of initial naphthalene (500 mg L^-1^) was utilized by immobilized *Pseudomonas* sp. consortium after 72 h of incubation. Further, initial naphthalene tolerance by immobilized *Pseudomonas* sp. consortium was highest (1475 mg L^-1^) with maximum specific degradation of 0.707 h^-1^. The gene sequence analysis of the naphthalene 1, 2-dioxygenase suggests a significant evolutionary distance among different microbial species and in few cases, intra-species variation was observed. A common mutation prone amino acid stretch inside Chain A of all three natural mutant variants of naphthalene 1, 2-dioxygenase were found at close proximity of the active site. Further, the rigid-flexible molecular docking showed better binging free energy of the mutant variant encoded by *Pseudomonas putida* strain KD10 than that of wild-type variant of naphthalene 1,2-dioxygenase. This common mutation prone amino acid stretch could aid the rational approaches of enzyme redesigning. Overall, this study summarises the application of bacterial cell immobilization in calcium alginate beads and development of the microbial consortium together for enhanced naphthalene biodegradation. However, further studies are required for the systematic optimization of naphthalene biodegradation in a real environment.

## Authors Contributions

KD and CG conceive the main hypothesis. KD design, performed all experiments, and wrote the manuscript. SS, IK, SB, DJ assisted KD in some experiments. MK, TM, PR, KCG performed the statistical analysis and wrote the manuscript. CG critically proofread and wrote the manuscript. All authors read the manuscript.

## Funding

University Grant Commission (UGC) Govt. of India, New Delhi, India (Grant No. VU/Innovative/Sc/17/2015).

## Acknowledgement

University Grant Commission (UGC) Govt. of India, New Delhi, India (Grant No. VU/Innovative/Sc/17/2015) is acknowledged by C.G. K.D. acknowledges Council of Scientific and Industrial Research (CSIR), Govt. of India, New Delhi, India for Senior Research Fellowship (SRF) sanction letter no. 09/599 (0082) 2K19 EMR-Z dated: 29/03/2019.

## Conflict of Interest

The authors declare that they have no conflict of interests.

## Supplementary section

**Table S1.** Nucleotide sequences used as primer in polymerase chain reactions

**Table S2.** Parameters of substrate inhibition kinetic model of naphthalene biodegradation by different *Pseudomonas putida* strains immobilized in calcium alginate beads.

**Table S3.** Enumeration of viable cell in calcium alginate beads.

**Table S4.** Gompertz’s growth curve model fit of *Pseudomonas putida* strain KD10 and *Pseudomonas* sp. consortium.

**Table S5.** Major amino acid residues of mutant variant of naphthalene 1, 2-dioxygenase _I250, V256_ involve in hydrophobic interaction with different ligands.

**Figure S1.** Colony morphology and catechol confirmatory test. **A.** Colony morphology of *Pseudomonas putida* strain KD10 after 24 h of incubation. **B.** Colony morphology of *Pseudomonas putida* strain KD10 after one week of incubation.

**Figure S2.** Molecular phylogenetic analysis of *Pseudomonas putida* strain KD10.

**Figure S3.** Effect of cell adaptation on naphthalene biodegradation

**Figure S4.** Mass spectra of the major metabolites detected during biodegradation of naphthalene in CSM as sole source of carbon and energy by *Pseudomonas putida* KD10.

**Figure S5.** Agarose gel electrophoresis of PCR products of catechol 2, 3-dioxygenase (*nah*H) gene and solvent efflux pump (*srp*ABC) system. **A.** Catechol 2, 3-dioxygenase (*nah*H). Lane 1 and 2 *nah*H pcr product of *Pseudomonas putida* strain KD9 and *Pseudomonas putida* strain KD10. **B.** Solvent efflux pump (*srp*ABC) system. Lane 1: *Pseudomonas aeruginoasa* ATCC 9027, Lane 2: *Pseudomonas putida* strain KD10. Lane M: 100bp DNA ladder (HiMedia, India), Lane 3: Negative control.

**Figure S6.** Nucleotide sequence chromatogram of the mutant naphthalene 1, 2-dioxygenase (*nah*Ac), (chain-A) encoded by *Pseudomonas putida* strain KD10.

