## Supplementary material for "Enhanced biodegradation of naphthalene by *Pseudomonas* sp. consortium immobilized in calcium alginate beads": Table S1

***Supplementary section***

| **Table S1.** Nucleotide sequences used as primer in polymerase chain reactions | | | | |
| --- | --- | --- | --- | --- |
| Gene | Nucleotide sequences | Annealing temperature | Product size | Reference |
| *nah*Ac | 5’-CCGCGGAAAACTTTGTGGGGGA-3’ | 57°C | 468 | Dutta el al. 2017 |
|  | 5’-GCCCAAACGTACGCTGAACCGA-3’ |  |  |  |
| *srp*A | 5’- CTGACCCTGACAACTGACTT-3’ |  | 455 | Dutta el al. 2018 |
|  | 5’- TCGACATAAATGGGGTCCAG-3’ |  |  |  |
| *srp*B | 5’- TCGAAACCAATTCCTGCAAC-3’ |  | 976 | Dutta el al. 2018 |
|  | 5’- CAGCGGAACAAAGAAGACAG-3’ |  |  |  |
| *srp*C | 5’- CTCGGATCGATAGCCAGTAC-3’ |  | 622 | Dutta el al. 2018 |
|  | 5’- AACAGTCCATCAAGATCGCT-3’ |  |  |  |
| *nah*H | 5'-TCACCATCCGGAAAAAGGCCGC-3' | 58°C | 200 | This study |
|  | 5'-AGATCGCCTTGCCCAGGTCCTT-3' |  |  |  |
