## Supplementary material for "Enhanced biodegradation of naphthalene by *Pseudomonas* sp. consortium immobilized in calcium alginate beads": Table S2

| **Table S2.** Parameters of substrate inhibition kinetic model of naphthalene biodegradation by different *Pseudomonas putida* strains immobilized in calcium alginate beads | | | | |
| --- | --- | --- | --- | --- |
| Strain | *q*_max_ (h^-l^) | *ks* (mg L^-1^) | *ki* (mg L^-1^) | *R*^2^ |
| A | 0.629 ± 0.04 | 304.7 ± 3.66 | 1033 ± 1.66 | 0.996 |
| B | 0.651 ± 0.19 | 322.4 ± 6.24 | 1234 ± 6.11 | 0.937 |
| C | 0.695 ± 0.24 | 309.2 ± 8.85 | 1324 ± 8.01 | 0.906 |
| D | 0.707 ± 0.23 | 298.8 ± 7.91 | 1475 ± 8.93 | 0.901 |
| Initial concentration of naphthalene = 500 -2500 mg L^-1^  A. *Pseudomonas putida* strain KD6, B*. Pseudomonas putida* strain KD9, C*. Pseudomonas putida* strain KD10. D. *Pseudomonas sp.* consortium. Each figure of the table represents the mean of three replicates. | | | | |
