## Supplementary material for "Enhanced biodegradation of naphthalene by *Pseudomonas* sp. consortium immobilized in calcium alginate beads": Table S3

| **Table S3.** Enumeration of viable cell in calcium alginate beads | | | |
| --- | --- | --- | --- |
| Strain | CFU gm^-1*^ | | |
|  | Day1 | Day 7 | Day 21 |
| A | 2.4×10^8^ | 2.16×10^8^ | 1.98×10^8^ |
| B |  | 2.01×10^8^ | 1.91×10^8^ |
| C |  | 2.14×10^8^ | 1.92×10^8^ |
| D |  | 2.19×10^8^ | 2.08×10^8^ |
| A. *Pseudomonas putida* KD 6, B. *Pseudomonas putida* KD9, C. *Pseudomonas putida* KD10, D. *Pseudomonas sp.* consortium. ^*^ Colony forming unit were measured from each gram of calcium alginate beads | | | |
