## Supplementary material for "Enhanced biodegradation of naphthalene by *Pseudomonas* sp. consortium immobilized in calcium alginate beads": Table S4

| **Table S4.** Gompertz’s growth curve model fit of *Pseudomonas putida* strain KD10 and *Pseudomonas sp.* consortium | | | | |
| --- | --- | --- | --- | --- |
| Strains | *a* | *xc* | *k* | *R^2^* |
| A. | 0.254 ± 0.00 | 3.65 ± 0.16 | 1.415 ± 0.39 | 0.964 |
| B. | 0.276 ± 0.01 | 3.66 ± 0.17 | 1.327 ± 0.37 | 0.960 |
| C. | 0.492 ± 0.01 | 3.23 ± 0.15 | 1.180 ± 0.28 | 0.971 |
| D. | 0.511 ± 0.01 | 3.26 ± 0.17 | 1.226 ± 0.34 | 0.960 |
| A. *Pseudomonas putida* KD10, B. *Pseudomonas sp.* consortium. C. *Pseudomonas putida* KD10 with 0.5gm% sucrose supplementation. D. *Pseudomonas sp.* consortium with 0.5gm% sucrose supplementation. | | | | |
