## Supplementary material for "Enhanced biodegradation of naphthalene by *Pseudomonas* sp. consortium immobilized in calcium alginate beads": Table S5

| **Table S5.** Major amino acid residues of mutant variant of naphthalene 1, 2-dioxygenase _I250, V256_ involve in hydrophobic interaction with different ligands | | | |
| --- | --- | --- | --- |
| Residues | Distance from the ligand (Å) | | |
|  | Naphthalene | Phenanthrene | Anthracene |
| Val 209 | 3.49 | 3.89, 3.87 | - |
| Phe 352 | 3.35, 3.21 | 3.43 | 3.87 |
| Phe 224 | - | 3.59, 3.98 | 3.48, 3.33 |
| Leu 307 | - | 3.65 | 3.76, 3.81 |
|  | π- staking bond distance (Å) | | |
| Phe 202 | 5.04, 5.14 | - | - |
| His 208 | 4.57 | - | 5.16 |
|  | π- staking bond angle (Degree) | | |
| Phe 202 | 67.76 | - | - |
| Phe 202 | 67.61 | - | - |
| His 208 | 80.58 | - | 77.19 |
