## Supplementary figures and images for "Enhanced biodegradation of naphthalene by *Pseudomonas* sp. consortium immobilized in calcium alginate beads"

### Figure S1

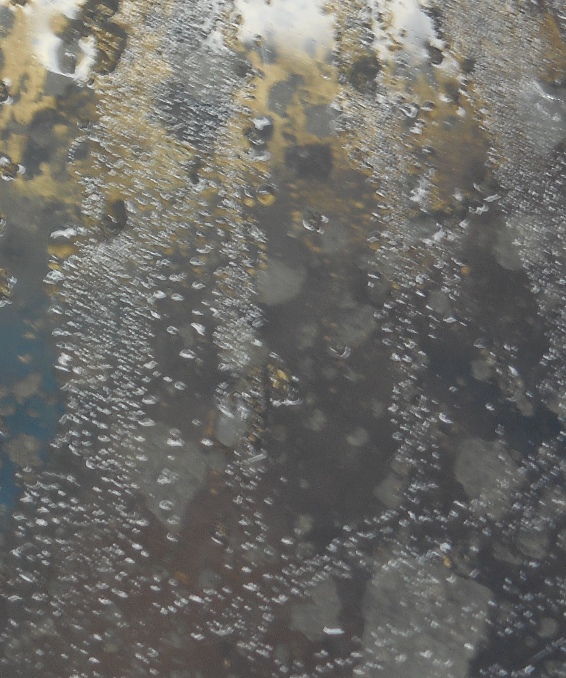

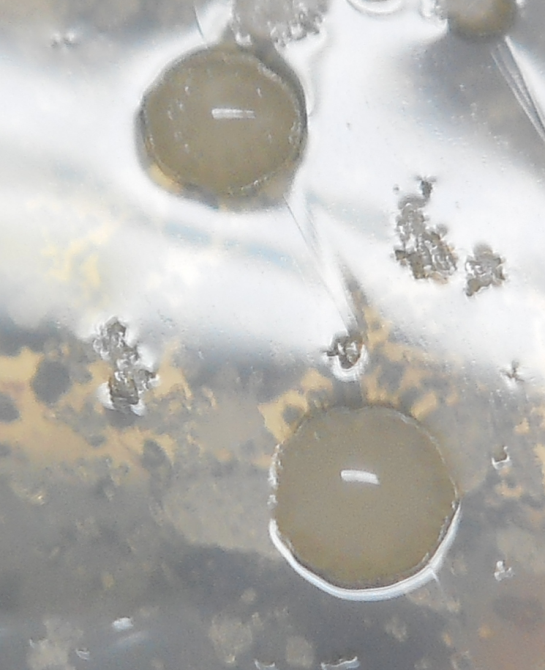

**B.**

**A.**

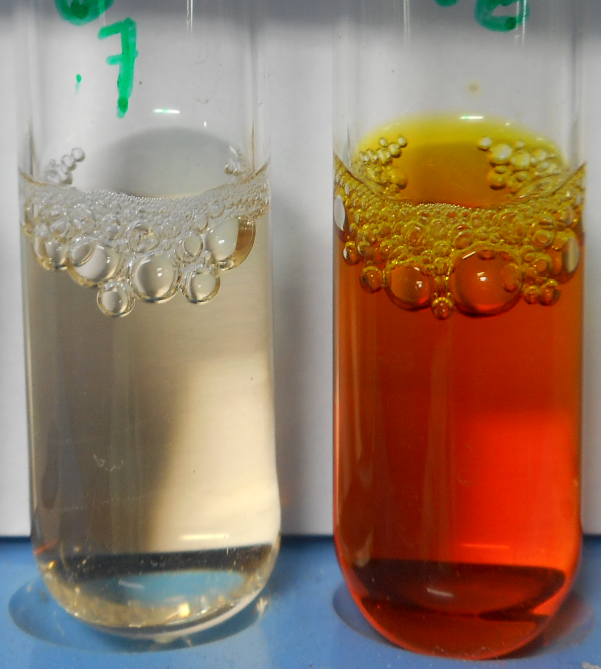

**(i)**

**(ii)**

**C.**

**Figure S1.**

### Figure S2

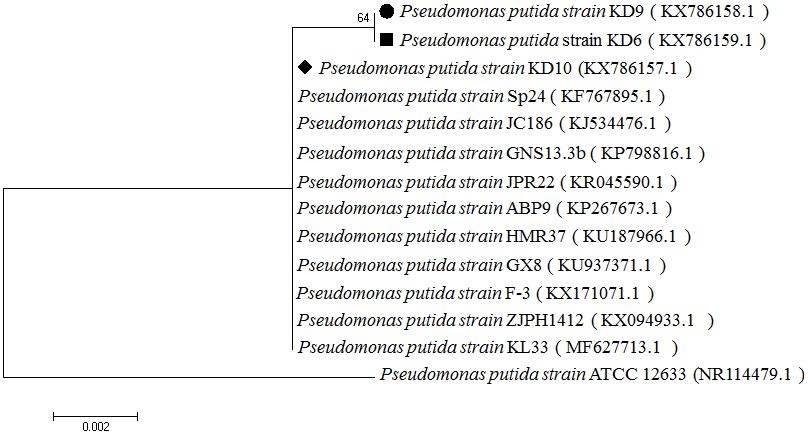

**Figure S2.**

### Figure S3

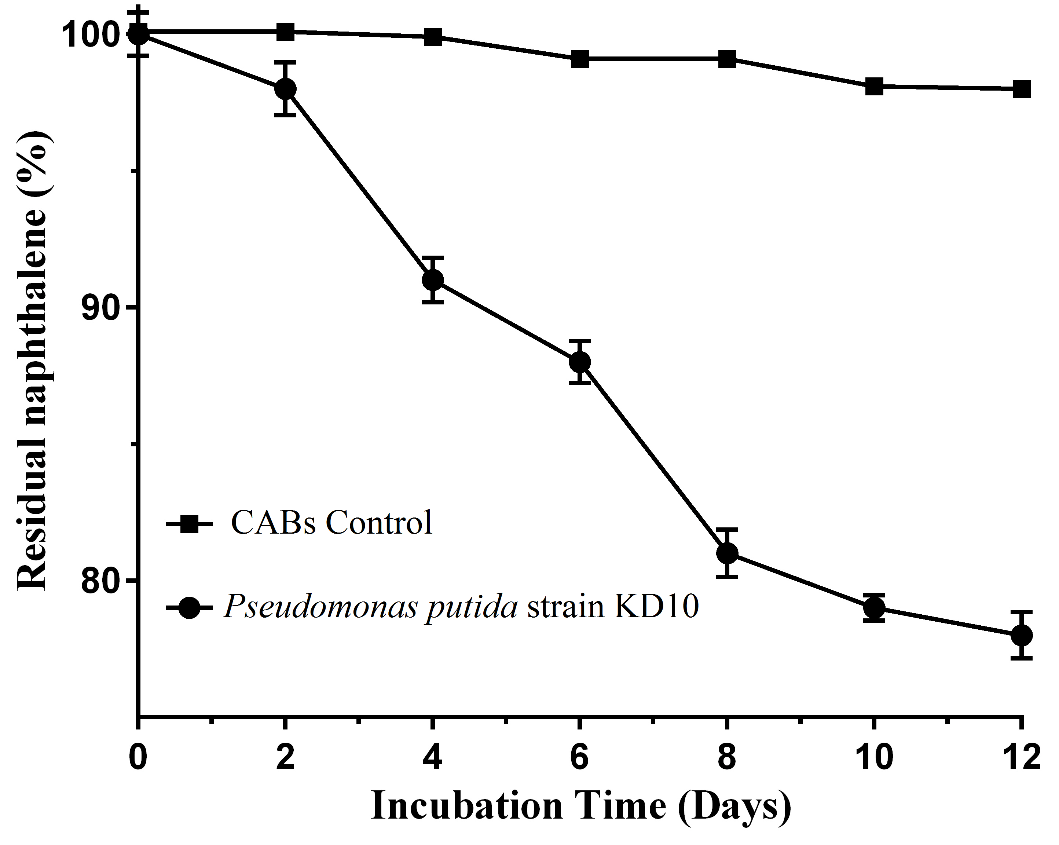

**Figure S3.**

### Figure S4

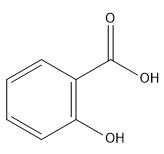

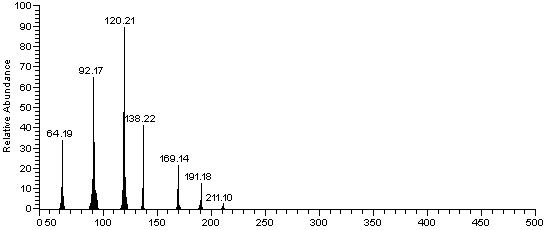

Salicylic acid

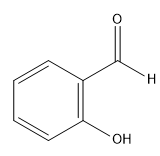

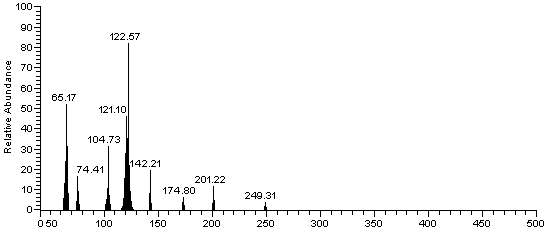

Salicylaldehyde

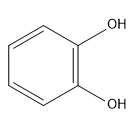

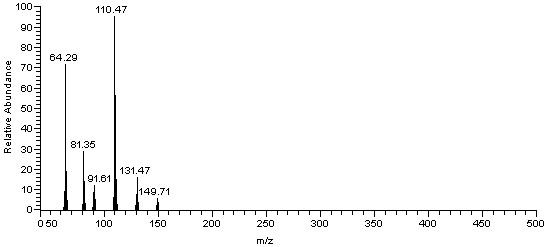

Catechol

Catechol

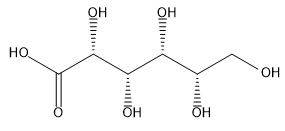

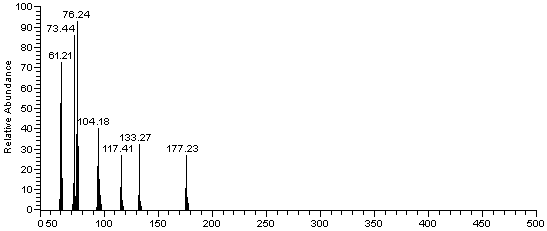

D-gluconic acid

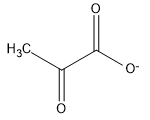

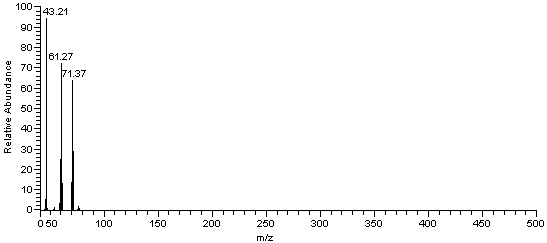

Pyruvic acid

**Figure S4.**

### Figure S5

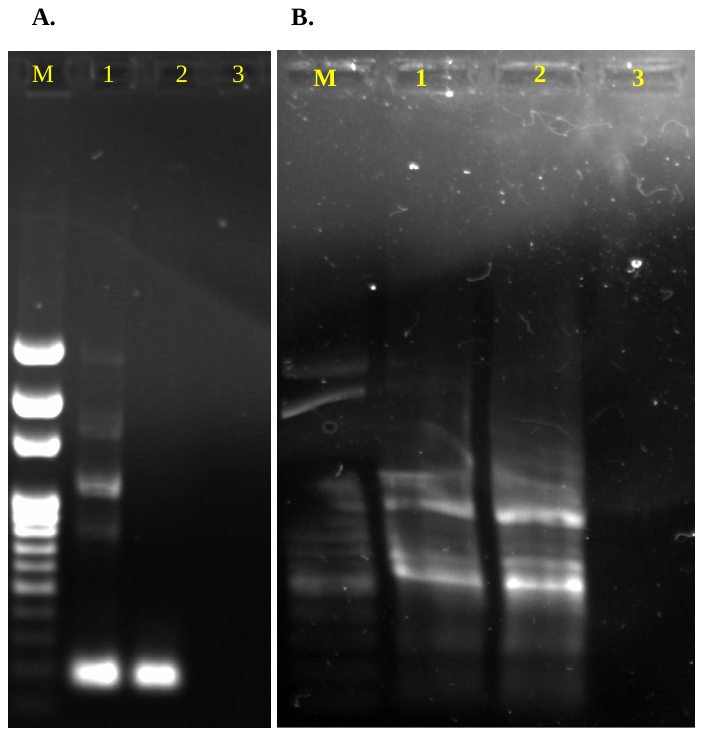

**Figure S5.**

### Figure S6

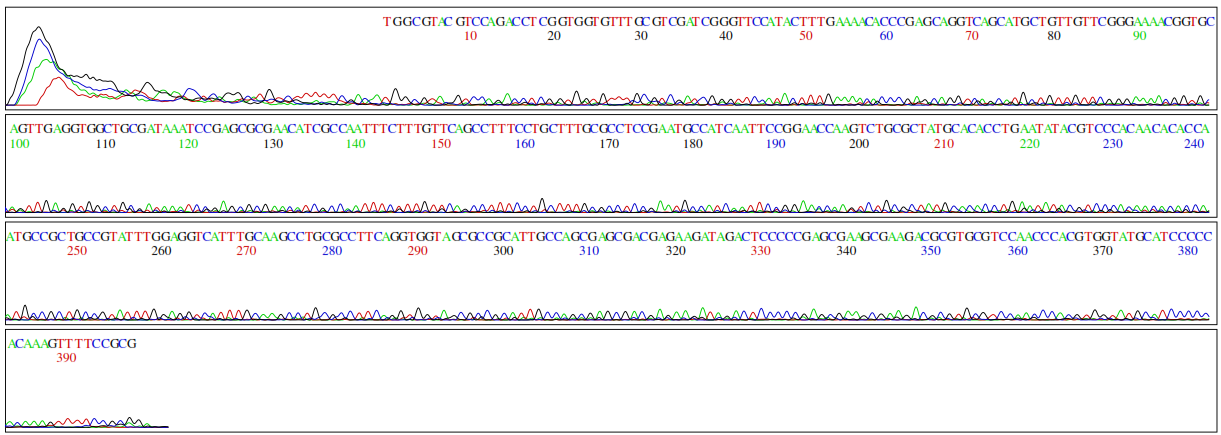

**Figure S6.**
